# High-dimensional spectral flow cytometry of activation and phagocytosis by peripheral human polymorphonuclear leukocytes

**DOI:** 10.1101/2024.12.01.626241

**Authors:** Evan R. Lamb, Ian J. Glomski, Taylor A. Harper, Michael D. Solga, Alison K. Criss

## Abstract

Polymorphonuclear lymphocytes (PMNs) are terminally differentiated phagocytes with pivotal roles in infection, inflammation, tissue injury, and resolution. PMNs can display a breadth of responses to diverse endogenous and exogenous stimuli, making understanding of these innate immune responders vital yet challenging to achieve. Here, we report a 22-color spectral flow cytometry panel to profile primary human PMNs on population and single cell levels for surface marker expression of activation, degranulation, phagocytosis, migration, chemotaxis, and interaction with fluorescently labeled cargo. We demonstrate the surface protein response of PMNs to phorbol ester stimulation compared to untreated controls in an adherent PMN model with additional analysis of intra- and inter-subject variability. PMNs challenged with the Gram-negative bacterial pathogen *Neisseria gonorrhoeae* revealed infectious dose-dependent changes in surface marker expression in bulk, population-level analysis. Imaging flow cytometry complemented spectral cytometry, demonstrating that fluorescence signal from labeled bacteria corresponded with bacterial burden on a per-cell basis. Spectral flow cytometry subsequently identified surface markers which varied with direct PMN-bacterium association as well as those which varied in the presence of bacteria but without phagocytosis. This spectral panel protocol highlights best practices for efficient customization and is compatible with downstream approaches such as spectral cell sorting and single-cell RNA-sequencing for applicability to diverse research questions in the field of PMN biology.

**Summary Sentence:** Here we report a 22-color spectral flow cytometry panel to profile primary human PMNs for markers of activation, degranulation, phagocytosis, migration, and chemotaxis using phorbol ester stimulation and bacterial challenge as proofs-of-concept.

## Introduction

Polymorphonuclear leukocytes (PMNs) are principal cellular responders to infection and inflammation in vertebrates. The granulocytic PMN population is predominantly composed of neutrophils, professional phagocytes with many specialized antimicrobial properties. Among these antimicrobial mechanisms are phagocytosis with subsequent phagolysosome maturation, active chemotactic migration toward pathogens, coordinated exocytosis of antimicrobial-containing granule subsets, and reactive oxygen species (ROS) generation.^1–4^ PMNs sense a variety of host- and pathogen-derived stimuli from the inflammatory milieu and integrate the resulting signals to coordinate their activation states and responses.^5–7^ Activation includes mobilization of surface proteins required for transmigration from the circulation to the site of infection/inflammation, downregulation of other surface proteins throughout this process through endocytosis or ectodomain shedding, and coordinated upregulation of proteins representative of primed antimicrobial activity.^8–14^ The breadth of stimuli and potential PMN reactions imply that PMNs have heterogenous responses to infection and injury. Approaches to measure the heterogeneity of PMN activation and responses on both single cell and population levels can elucidate how PMNs respond in a variety of conditions.^15–19^

A time-tested methodology to characterize and quantify leukocyte activation in response to diverse stimuli is flow cytometric analysis. Flow cytometry has proven to be an advantageous approach given its increased availability, high throughput nature, relatively small cell numbers needed, easy differentiation of surface versus intracellular expression, and population versus single cell analytics, among other positive attributes.^20–23^ Among flow cytometric methods, conventional flow cytometry is limited by the number of markers that can be analyzed in a sample at a given time due to technical constraints which prevents integration of large numbers of parameters on single cells at the same time to make direct comparisons. Partially as a result from such limitations, many flow studies on PMN biology have analyzed markers as single-stained samples focused on granulocyte development/ontogeny^24–26^, or select parameters that reflect PMN capacity for migration/chemotaxis, phagocytosis, ROS generation, NETosis^27^, or antimicrobial release in isolation.^14,28–31^ To surmount limitations presented by conventional flow cytometry, technologies have been engineered that expand high-dimensional flow cytometric capacity. One such high-dimensional method is cytometry by time of flight (CyTOF) which uses heavy metal-conjugated antibodies to label cells of interest and identify target positivity and expression levels. This technology enables analysis of vast markers in a single sample and has been successfully applied to PMNs.^32,33^ However, CyTOF is a costly method and requires sample destruction for data generation.^34,35^

Another advanced flow cytometric methodology is spectral flow cytometry in which the full fluorescence spectrum of individual antibody-conjugated fluorochromes can be collected from each excitation laser in a cytometer’s configuration, allowing for a full ‘spectral fingerprint’ to be collected and identified. This permits many more fluorochrome combinations than conventional flow cytometry, and thereby more cell markers, to be examined in a single experimental condition.^36,37^ Spectral flow cytometry has been effectively deployed to analyze human PMN activation and identify subsets in healthy and diseased states by examining up to 15 surface markers at once.^22,38–41^ Recent advances in spectral flow cytometry technology, including spectral cell sorting and single-cell analytical software, enable the rapid and insightful analysis of high-dimensional datasets.^42,43^

With the above pros and cons in mind, we sought to design a flow cytometry panel that 1) analyzes mature human PMNs, 2) does so in a high-dimensional manner, 3) does so without sample destruction to allow for downstream sorting/analysis, 4) focuses on both PMN functionality and activation, and 5) is adaptable to diverse research questions in the field. Such a methodology could advance the understanding of PMN activation and diversity as well as the contribution that such diversity provides to outcomes in inflammation, infection, and injury.

Here, we present a 22-color spectral flow cytometry panel to profile the activation of mature human PMNs in response to diverse stimuli. The panel described here was designed with particular emphasis on PMN (opsono)phagocytic receptors, degranulation markers, migratory proteins, chemokine receptors, and the option to fluorescently-label cargo, such as microbes.^7^ We describe best practices for using the panel in different laboratories. We demonstrate that the panel can identify PMNs in a population that respond to phorbol ester stimulation and infection by the bacterial pathogen *Neisseria gonorrhoeae*. The panel can be customized for different fluorochrome-marker pairings, and common fluorochromes/channels are left available for incorporation of other markers of interest, enabling its adaptation to many research endeavors.

## Materials and Methods

### PMN Isolation from human subjects

Human subjects research was conducted in accordance with the University of Virginia Institutional Review Board for Health Sciences Research under protocol #13909. Informed and written consent was obtained from each human subject. Primary human PMNs were collected via venipuncture from peripheral blood of healthy human donors in accordance with institutional Human Subjects in Research guidelines as previously described by Ragland and Criss.^44^ Briefly, venous blood was collected into heparin-coated vacutainer tubes and fractionated by dextran sedimentation to enrich for leukocytes. Granulocytes were further purified by Ficoll-Paque^TM^ density centrifugation with DPBS (Gibco) + 0.1% glucose (Ricca Chemical; DPBSG). The Ficoll-PBS interface enriched for monocytes and depleted of granulocytes was collected for CD14 gate setting. The granulocyte pellet was then resuspended, lysed with endotoxin-free water to remove remaining erythrocytes, and resuspended in DPBSG on ice and enumerated using a hemacytometer.

### Neisseria gonorrhoeae Growth and Labeling

An FA1090^45^ strain of *N. gonorrhoeae* which constitutively expresses the PMN-binding surface protein OpaD and no other opacity-associated proteins^46^ was streaked on gonococcal base media agar plates and incubated at 37°C with 5% supplemental CO_2_ for 16 hours.^47,48^ Single colonies were then swabbed into Hank’s balanced salt solution (HBSS, with 1.2mM calcium and 1mM magnesium, Gibco) with 10mM HEPES (Sigma-Aldrich) pH 7.4 and 5mM sodium bicarbonate (HBSS+) to a concentration of 1.5e8 bacteria per mL and labeled with CellTrace Blue (ThermoFisher) for 20min at 37°C. Bacteria were then pelleted and resuspended in HBSS + 2% BSA to quench remaining CellTrace Blue dye. Un-labeled bacteria were used as non-fluorescent controls.

### PMN Adherence and Stimulation

To simulate post-migration status of innate immune cells at inflamed mucosa, isolated primary human PMNs were primed with 10nM recombinant human interleukin-8 (IL-8, R&D Systems) in Roswell Park Memorial Institute Medium (RPMI, Gibco) + 10% (v/v) heat-inactivated fetal bovine serum (Hyclone, RPMI + 10% FBS) as in Ragland and Criss.^44^ PMNs were then allowed to settle and adhere onto 25mm plastic cover slips (Sarstedt) in 6-well tissue culture plates in 1mL medium at 37°C, 5% CO_2_ for 30-60min. Following adherence, PMNs were either left untreated, stimulated with 10ng/mL phorbol 12-myristate 13-acetate (PMA) (Sigma-Alrich), or infected with *N. gonorrhoeae* at a multiplicity of infection of either 1 or 10 bacteria per PMN for 60min at 37°C, 5% CO_2_. Controls included unstained/untreated samples and single fluorochrome samples. Each condition consisted of two wells in a 6-well plate with 2e6 PMNs per well which were pooled following stimulation.

### PMN Washing and Labeling

Following 60min of stimulation/infection, 500mM EDTA (Sigma-Aldrich) was added to adhered PMNs to a final concentration of 0.5mM. PMNs were gently resuspended using a cell scraper (Falcon). Cells from two wells per condition were pooled into a 15mL conical tube and centrifuged at 1900 x g for 7min at 4°C. Medium was removed via aspiration to approximately 50μL and PMNs were gently resuspended in 2mL ice cold HBSS+ and washed likewise twice more. Following the final wash, PMNs were resuspended in 200μL ice cold HBSS+ and transferred to a V-bottom 96-well plate on ice. Additional samples added to the 96-well plate included: PMNs which had been untreated and left in suspension to be used both for full antibody staining and for unstained controls; suspension PMNs on ice mixed 1:1 with suspension PMNs which were heat-killed (65°C for 5min) for viability gate setting; and granulocyte-depleted DPBSG-Ficoll interface ‘buffy-coat’ enriched with monocytes for CD14 dump gate setting. The plate was centrifuged at 1900 x g for 7min at 4°C, 100μL were removed from each well via multichannel pipet, and a 1:1000 dilution of Zombie Near InfraRed (ZNIR) Live-Dead dye (BioLegend) was added to the full-stain and single/gate-setting stain wells per manufacturer’s directions (15min at room temperature in the dark) and pellets gently resuspended. 100μL of Flow Staining Buffer (eBiosciences) was then added to each well to quench ZNIR dyes. The plate was centrifuged as above, 150μL were removed from each well via multichannel pipet, and 150μL Flow Staining Buffer was added. 150μL was removed from each well via multichannel pipet so that each well contained 50μL of Flow Staining Buffer and pelleted cells. Flow Staining Buffer was added to wells followed by individual antibodies as indicated in Table 1 to a total of 100μL per well. Cells were gently resuspended with staining buffer/antibody mixtures and incubated at 4°C for 30min in the dark. The plate was then centrifuged as above, 50μL was removed from each well, and pellets were gently washed three times in sterile PBS. The final wash was into a final volume of 100μL of PBS + 1% (v/v) paraformaldehyde (Electron Microscopy Sciences). The plate with fixed samples was stored at 4°C in the dark wrapped in aluminum foil for no more than three days before analysis on the spectral flow cytometer.

**Table 1.**
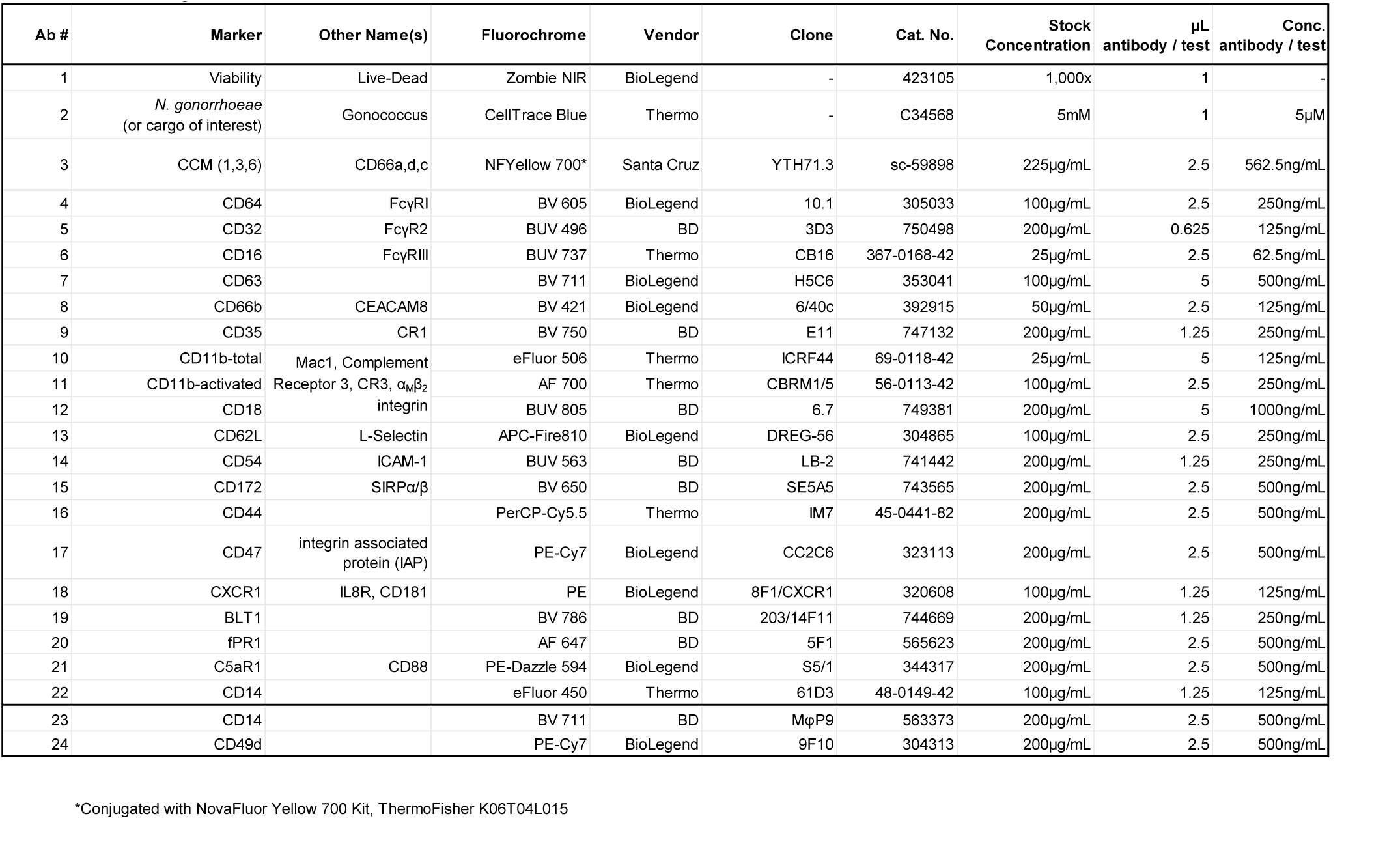
Antibodies and Reagents.

### Spectral Flow Cytometry Acquisition

Fixed samples were run on a Cytek Aurora spectral flow cytometer with a 20mW 355nm, 50mW 488nm, 100mW 405nm, 50mW 561nm, and 80mW 640nm 5-laser configuration. Samples were run in a V-bottom 96-well plate using the autosampler apparatus within three days after fixation. Unmixing and Spillover correction was performed in SpectroFlo (Cytek) software.

### Antibody-fluorochrome conjugation and titration

All fluorescently labeled antibodies were obtained from commercial suppliers (Table 1), with the exception of the anti-CEACAM 1,3,6 antibody. The anti-CEACAM 1,3,6 antibody was labelled with NovaFluorYellow 700 using the NovaFluor Antibody Conjugation Kit (ThermoFisher) and conjugated per manufacturer’s protocols. Each fluorescently labelled antibody was titrated to establish the lowest concentration that maximized the fluorescence intensity differential between labelled and unlabeled cells. The highest concentration of antibody used was based on manufacturer suggestions (typically 5 µl) and serially diluted to 0.5x, 0.25x, and 0.125x final concentrations. The fluorescence intensity ratio for each antibody concentration relative to unstained control cells was established and the lowest antibody concentration that provided the largest ratio of fluorescence intensity of positive cells to negative cells was used in subsequent assays.

### Imaging flow cytometry analysis of N. gonorrhoeae infected PMNs

Primary human PMNs were isolated and infected as described above with *N. gonorrhoeae*. Bacteria had been labeled with both CellTrace Blue and CellTrace Yellow (ThermoFisher) per manufacturer’s protocols to be detected on both the Cytek Aurora spectral flow cytometer (both fluorochromes) and the Cytek ImageStream X MkII imaging flow cytometer (CellTrace Yellow) with single stained and unstained bacteria as controls. Following infection, cells were collected and fixed as above, and data collected on each cytometer with appropriate single stained controls. PMNs were assayed via imaging flow cytometry at 60x magnification using brightfield to collect micrographs of individual cells, side scatter channels, and the 561nm (100mW) excitation laser to collect CellTrace Yellow-Gc median fluorescence intensity. Ten-thousand individual, focused singlet PMN events were collected for each sample and data was analyzed using the IDEAS 6.2^®^ software package.^48,49^

### PMN Cell Sorting

PMNs were infected with CellTrace Yellow and CellTrace Blue dual labeled *N. gonorrhoeae* as described above, scraped from coverslips, stained with Zombie NIR, and run on a Cytek CS spectral Cell Sorter flow cytometer. Live unmixing was performed and PMNs were sorted into CellTrace Blue negative, low and high subgated populations as well as a non-gated group into Eppendorf tubes. These sorted PMNs were subsequently run a Cytek Aurora spectral flow cytometer to assess post-sorting viability via Zombie NIR exclusion using a one-to-one mixture of viable and heat-killed PMNs as a control.

- *Statistics, Analyses, and Data Availability:* Statistical analyses were performed as indicated in the figure legend for intra donor variability of individual markers within the PMN flow cytometry panel. Data were analyzed and prepared using SpectroFlo (Cytek), FCS Express (De Novo), IDEAS (Amnis), and GraphPad Prism software. High-dimensional analyses were performed on spectral flow cytometry data which was unmixed in SpectroFlo software prior to uploading into OMIQ cloud flow cytometry software for uniform manifold approximation and projection (UMAP) dimensional reduction processing and analysis.^42^ Data was cleaned using FlowAI within OMIQ, gated onto viable single PMN events, and UMAP analyses run on adherent PMN data files with or without PMA stimulation with all collected fluorescent features except those used for gating or CellTrace Blue. OMIQ’s default UMAP settings were used. Raw flow cytometry data is available through the MyFlowCyt flow cytometry repository under the Experiment title: PMN Spectral Flow Panel, and Experiment ID: (FR-FCM-Z8EX). http://flowrepository.org/experiments/8664.

## Results

### Spectral Flow Cytometry Panel Design for PMN Function and Activation

We designed the following spectral flow cytometry panel for mature human PMN functions (Fig. 1). Analyses were performed on human peripheral blood PMNs, which were freshly isolated on the day of each experiment from healthy subjects as described in Materials and Methods.^44^

**Figure 1.**
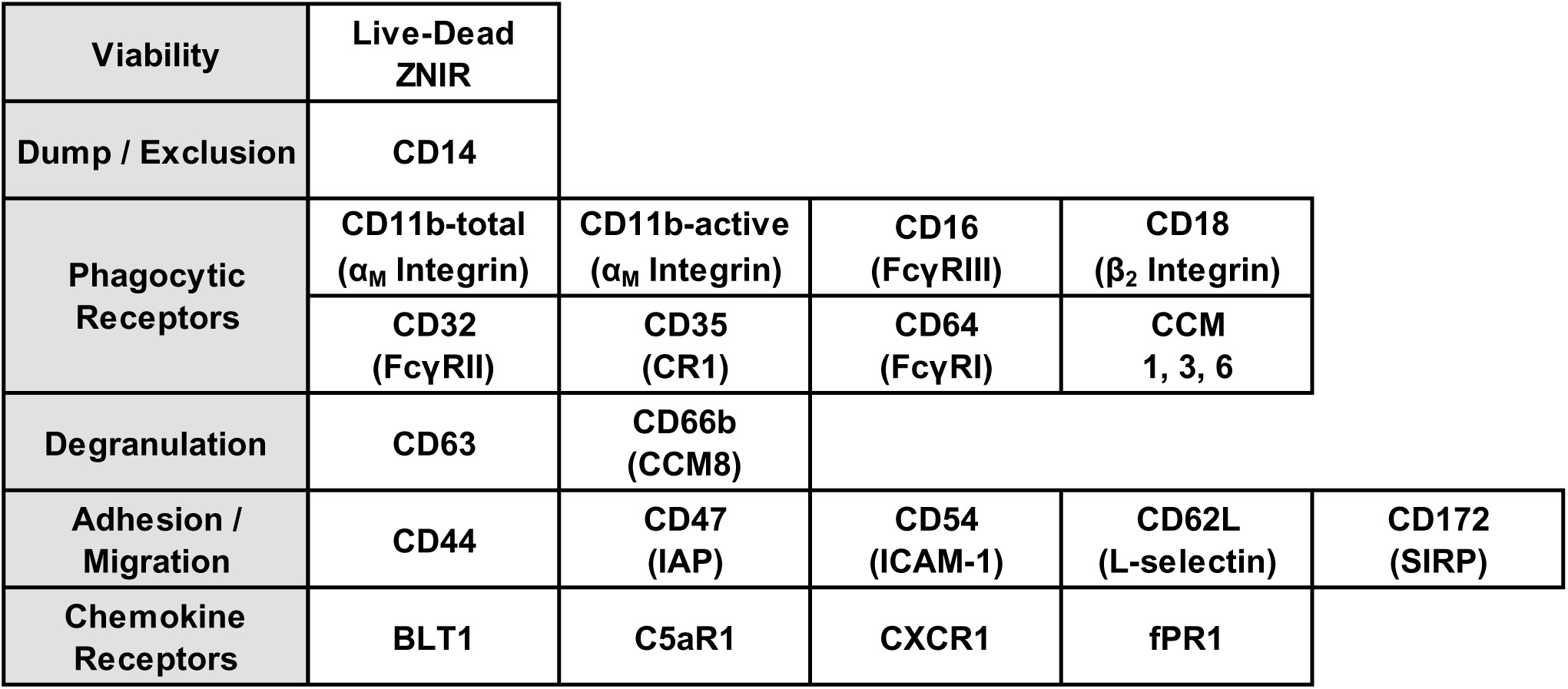
PMN Spectral Flow Cytometry Panel Surface Markers/Parameters Grouped by Functional Category. Schematic of the selected parameters in the PMN spectral flow cytometry panel, organized by function. Where applicable, cluster of differentiation (CD) label is listed with other common names in parentheses. Zombie Near-Infrared (ZNIR); FcγReceptor (FcγR); Complement Receptor (CR); Carcinoembryonic antigen-related cell adhesion molecules (CCM); Integrin Associated Protein (IAP); Intercellular Adhesion Molecule 1 (ICAM-1); Signal-regulatory protein (SIRP).

#### Cell viability and exclusion of non-PMNs

This panel was designed with an amine-reactive live-dead stain to exclude non-viable cells (Zombie Near Infrared, ZNIR). The major co-purifying cell type in the PMN preparations are CD14^High^ monocytes; therefore, an anti-CD14 antibody was added to exclude CD14^High^ cells from downstream analysis.^21^

#### PMN phagocytosis

The panel contains antibodies against CD64 (FcγR1), CD32 (FcγRII), and CD16 (FcγRIII), which mediate phagocytosis of IgG-opsonized cargo^50^, and antibodies against CD35 (complement receptor 1, CR1) and CD11b and CD18 (complement receptor 3, CR3) for complement C3b-, iC3b-, and C3d(g)-mediated opsonophagocytosis.^33^ CD11b undergoes activation-dependent conformational changes; thus antibodies against total and active forms of CD11b were included.^51^ We also included an antibody against human carcinoembryonic associated cellular adhesion molecules (CCMs), which serve as non-opsonic phagocytic receptors for many pathogens including *N. gonorrhoeae*.^52,53^ Of the CCM family members, the antibody used in this study recognizes the granulocyte-expressed CCMs -1, -3, and -6 (but not CCM8/CD66b; Table 1).

#### PMN degranulation

The panel contains antibodies against the primary/azurophilic granule protein CD63 and the secondary/specific granule protein CD66b. Both are well described markers for individual granule subsets that are sequentially exocytosed from PMNs upon activation.^9^

#### PMN migration and chemotaxis

The panel includes antibodies against CD62L (L-selectin) which is shed as PMNs migrate to target sites^11,12^, CD54 (ICAM-1), CD172 (signal regulatory protein, SIRP), CD44, and CD47 (integrin associated protein, IAP) which is also a ligand for CD172. These receptors enable nuanced, context- and location-specific migratory responses of immune cells in infection and inflammation.^54,55^ The panel also contains antibodies against chemotactic receptors that are known to promote directional migration and PMN activation: CXCR1 for IL-8, BLT1 for leukotriene-B4 (LTB_4_), fPR1 for formylated peptides, and C5aR1 for the anaphylatoxin C5a of the complement cascade.^56^

The selected markers serve as broad examples of PMN activation/stimulation and phenotypic functional groups. They also reflect underlying PMN biology of granule mobilization to the plasma membrane (ex: CD63, CD66b, CD11b, CD18)^9^, endocytic downregulation of surface markers to ablate signal reception (ex: fPR1, C5aR1)^10^, and ectodomain shedding (CD16, CD62L).^11–13^

### Fluorochrome selection and panel similarity and complexity

Cognate fluorochromes for each marker were selected to optimize the functionality of the spectral flow cytometry panel while maximizing the number of markers to include (Fig. 2A). Considerations incorporated into panel design included cytometer laser and detector array configuration, spectral overlap between fluorochromes, epitope densities, and brightness indexes of the fluorochromes. However, given the intrinsic variability of human PMNs and the range of activation-dependent surface expression, it was challenging to select appropriate fluorochrome-marker pairings, and all selections required testing and validation within the context of the broader panel. Where possible, markers known to be in the same subcellular (granule) location were paired with fluorochromes with distinct spectra to minimize spectral overlap increasing the overall resolution of the panel.^8,57^ While most fluorescent antibodies used here are commercially available, the anti-CCM antibody was conjugated in-house to NovaFluor Yellow 700. The final PMN panel pairings yielded similarity and complexity values which were calculated using Cytek’s Full Spectrum Viewer, with a lower overall complexity score being optimal (Fig. 2B).^57^

**Figure 2.**
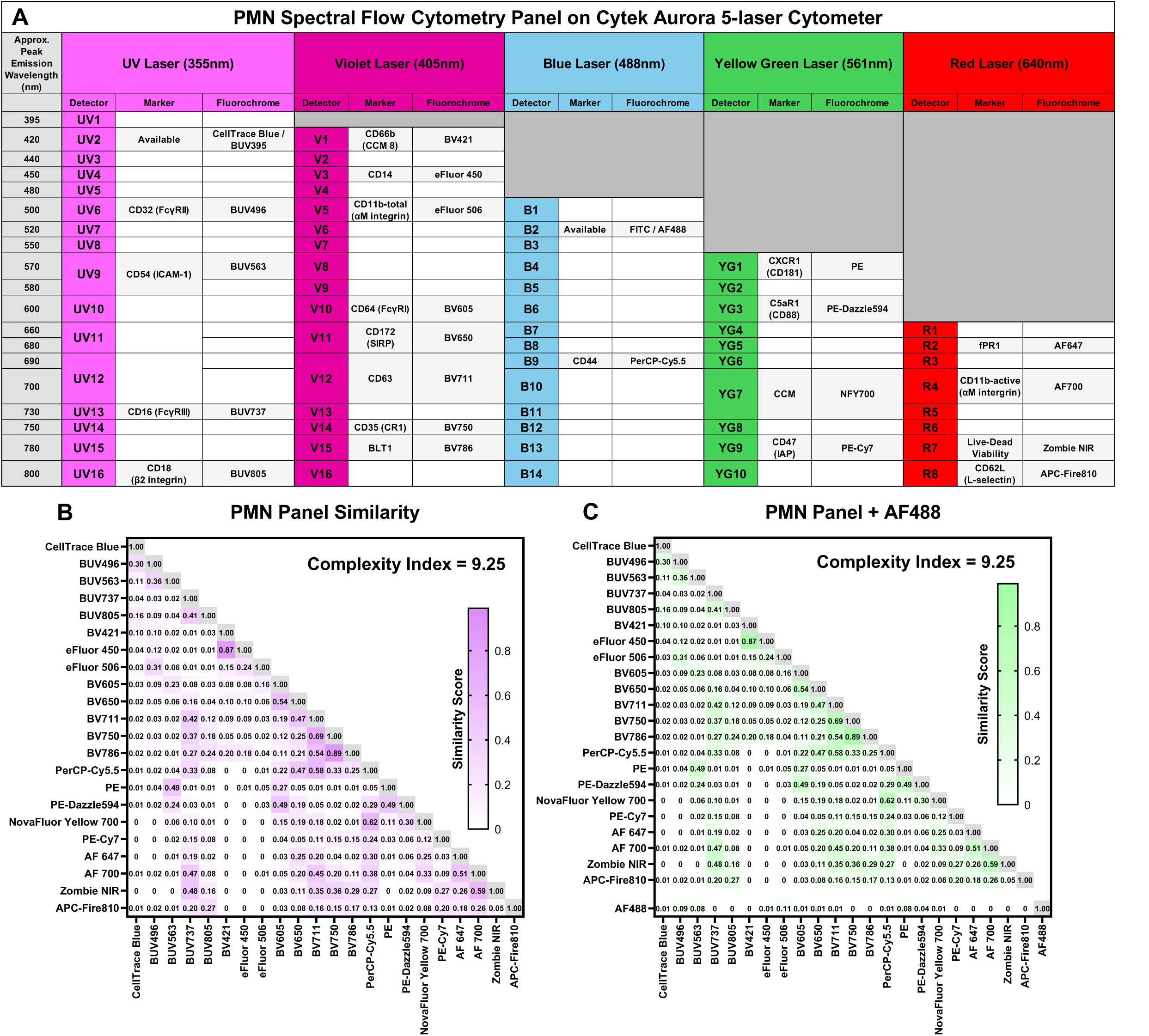
PMN Spectral Flow Cytometry Panel Surface Marker Fluorochrome Pairings, Similarity, and Complexity. **(A)** Representation of the Cytek Aurora 5-laser spectral flow cytometer detector array with 355-, 405-, 488-, 561-, and 640-nm laser configuration. Approximate peak emission wavelengths are listed top to bottom for each laser with corresponding detector array. Fluorochromes are listed in their peak detector slot with their cognate surface marker/parameter. The UV2 (CellTrace Blue/BUV395) and B2 detector channels have been left available for customization. **(B)** The similarity of each of the tested 22 fluorochromes’ predicted spectral fingerprints compared to each other was calculated using Cytek’s Full Spectrum Viewer. Calculated similarity is shown in each cell with a value of 0.0 indicating no similarity and 1.0 indicating exact similarity. The overall complexity index is shown at the top right. **(C)** The similarity and complexity are shown as in panel (B) but with the AlexaFluor 488 (AF488) fluorochrome included.

To maximize the adaptability of this panel to different research questions, the 355nm excitation/420nm peak emission (BUV395, CellTrace Blue; UV2 detector) and 488nm excitation/520nm peak emission (FITC, AlexaFluor488; B2 detector) channels were left available. Here, we used the BUV395 channel to pre-label *N. gonorrhoeae* with the amine-reactive dye CellTrace Blue. The FITC/AlexaFluor 488 (AF488) channel is a popular choice for many antibody conjugations and functional dyes and was left unused in this panel. However, addition of AF488 to the panel did not appreciably alter the calculated similarity matrix or complexity index of the panel (Fig. 2C).

### Panel gating strategy

Purified cells were first gated on events which passed FlowAI cleaning within OMIQ. These events were then gated into the PMN/granulocyte population based on characteristic forward and side scatter profiles (FSC-A, SSC-A; Fig. 3A i), and followed by selection of single cell events (Fig. 3A ii). Singlets were then sub-gated on CD14^Low^ events to exclude contaminating monocytes (Fig 3A iii).^21^ The CD14 gate was set using a mixture of the PMN preparation and the Ficoll-PBS interface (collected and saved on ice during the preparation process) which is enriched in CD14^High^ monocytes (Fig 3B). Finally, CD14^Low^ singlets were gated on live cells for subsequent analyses (Fig. 3A iv). The live-dead gate was set using a 1:1 mixture of cells that were killed by heating at 65°C for 5min and cells that were kept on ice before ZNIR staining (Fig. 3C).

**Figure 3.**
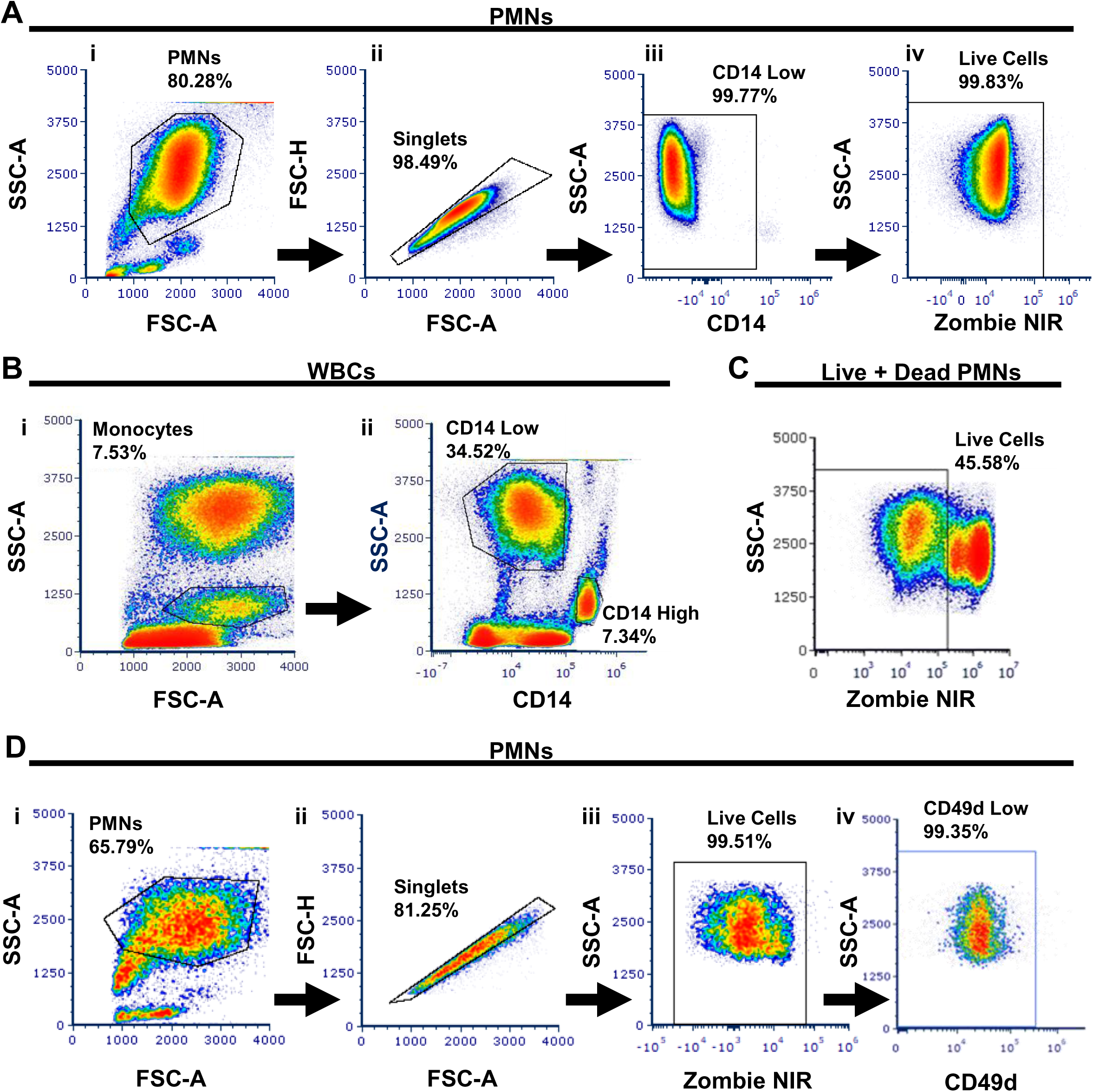
Gating Strategy for Live Primary Human PMNs. **(A)** The PMN/granulocyte population was gated from Ficoll-purified cells (see Materials and Methods) based on characteristic forward and side scatter profiles (i; FSC-A, SSC-A), followed by gating on singlet cells (ii), CD14 Low events (iii), and live cells (iv; Zombie near-infrared (NIR) exclusion). **(B)** The CD14 gate was set using a mixture of purified PMNs and cells collected from the Ficoll-PBS interface during PMN preparation which is enriched in CD14 High Monocytes. The Monocyte gate was set on the characteristic FSC-A and SSC-A profiles (i) with this gate being used to delineate CD14 High from CD14 Low populations (ii). **(C)** The viability (Zombie NIR) gate was set using a 1:1 mixture of viable and heat-killed PMNs. **(D)** The PMN/granulocyte population was gated on characteristic FSC-A and SSC-A profiles (i) followed by singlet cells (ii) and live cell events (iii) as described above. Live cells were gated into the CD49d Low population (iv) which was determined by labeling UltraComp Beads to determine neutrophil and eosinophil representation within the PMN gate.

While most PMNs in circulation are neutrophils, eosinophils also co-purify in the granulocyte preparations. Separate experiments were conducted in which live singlet PMNs were further gated on CD49d to discriminate eosinophils (CD49d+) from neutrophils (CD49d-; Fig 3D).^58,59^ PMN preparations were further evaluated for CD16, CD66b, and CD11b positivity to verify neutrophil predominance (Supplemental Fig. 1).^40^ These results showed the PMN preparations from healthy individuals to contain less than 1% eosinophils.^59^

### Analysis of intra- and intersubject variability in the PMN response to phorbol ester treatment

The panel was applied to adherent, IL-8 treated primary human PMNs^44^ under three conditions to model different types of stimulation: 1) PMNs with no further stimulus as a baseline, 2) PMNs that were also treated with phorbol myristate acetate (PMA), a potent protein kinase C agonist with known neutrophil-activating properties^60^, and 3) PMNs which were also infected with CellTrace Blue-labeled *N. gonorrhoeae*. The spectral flow cytometry panel was applied to measure variability in PMN responsiveness over time (>1 month between experiments), and to measure inter-subject variability for PMNs from three unrelated individuals. For inter-donor variability, three biological replicates of Subject #1 were combined and analyzed against the single replicates for the other two subjects.

For PMNs from the same subject, 10 of the 19 parameters increased in surface expression with PMA stimulation (CCM, CD64, CD63, CD66b, CD18, CD11b-total, CD11b-active, CD47, fPR1, and BLT1; Fig. 4A). Of the 19 markers, another 4 decreased in surface expression on PMNs treated with PMA (CD16, CD35, C5aR1, and CD62L; Fig. 4B). The remaining 5 markers showed no consistent trends between biological replicates (CD32, CD54, CD172, CD44, CXCR1; Fig. 4C).

**Figure 4.**
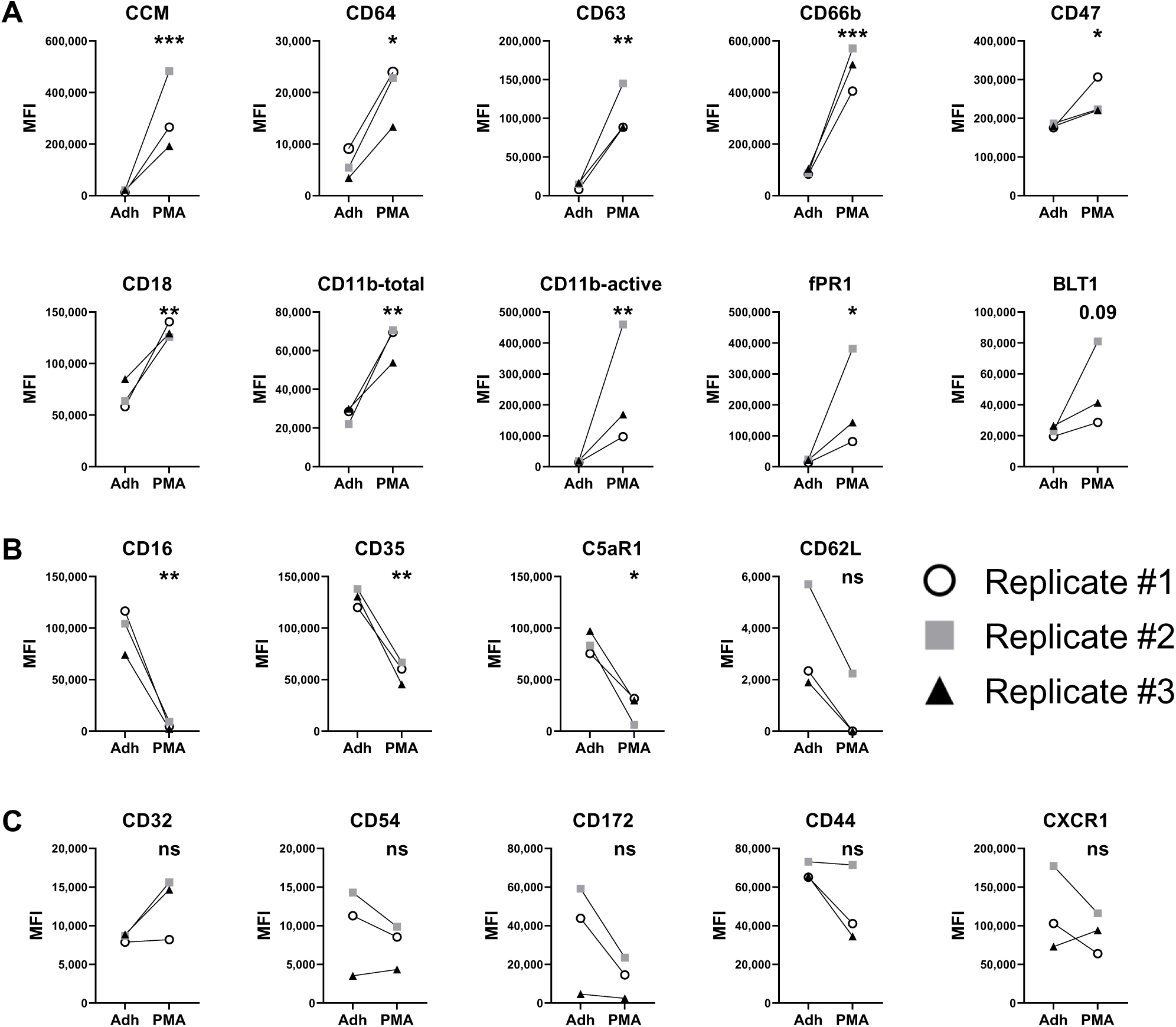
Assessing PMN Activation by Phorbol Ester and Intra-subject Variability Using High-dimensional Spectral Flow Cytometry. PMNs were purified from a single individual (Subject #1) on three separate days, represented by different colors/shapes. The median fluorescence intensity (MFI) was calculated for each parameter of the spectral flow cytometry panel. **(A)** Proteins with greater surface expression following PMA treatment compared to adherence alone (Adh) in each replicate. **(B)** Proteins with reduced surface expression following PMA treatment compared to adherence alone in all replicates. **(C)** Proteins with no consistent change in surface MFI between replicates. Statistical analyses were performed on unpaired log-transformed data, Student’s *t*-test. P-value indicated (0.05<p<0.10) or *=p<0.05, **=p<0.01, ***=p<0.001, ****=p<0.0001, ns = not significant. Negative MFI values at the population level were set to a value of 1.0 for graphing, log transformation, and subsequent statistical analyses.

For inter-subject responses, PMNs from three unrelated subjects showed consistently increased surface expression for 8 markers upon PMA stimulation (CCM, CD64, CD32, CD63, CD18, CD11b-active, CD47, and fPR1; Fig. 5A). For 3 other markers (CD66b, CD11b-total, and BLT1), PMA stimulation increased their surface expression for Subject #1 and #3’s PMNs whereas Subject #2’s PMNs did not appreciably change (Fig. 5B). Six markers decreased in all three subjects’ PMNs after PMA treatment (CD16, CD35, CD172, CD62L, CD44, and C5aR1; Fig. 5C). We note that two of these markers, CD172 and CD44, gave Subject #1 an overall decreased response when averaged, despite the response variability on each day (Fig. 4C). As seen for the replicates from Subject #1 (Fig. 4), the three unique subjects’ PMNs did not have consistent responses to PMA treatment in the surface expression of CD54 or CXCR1 (Fig. 5D).

**Figure 5.**
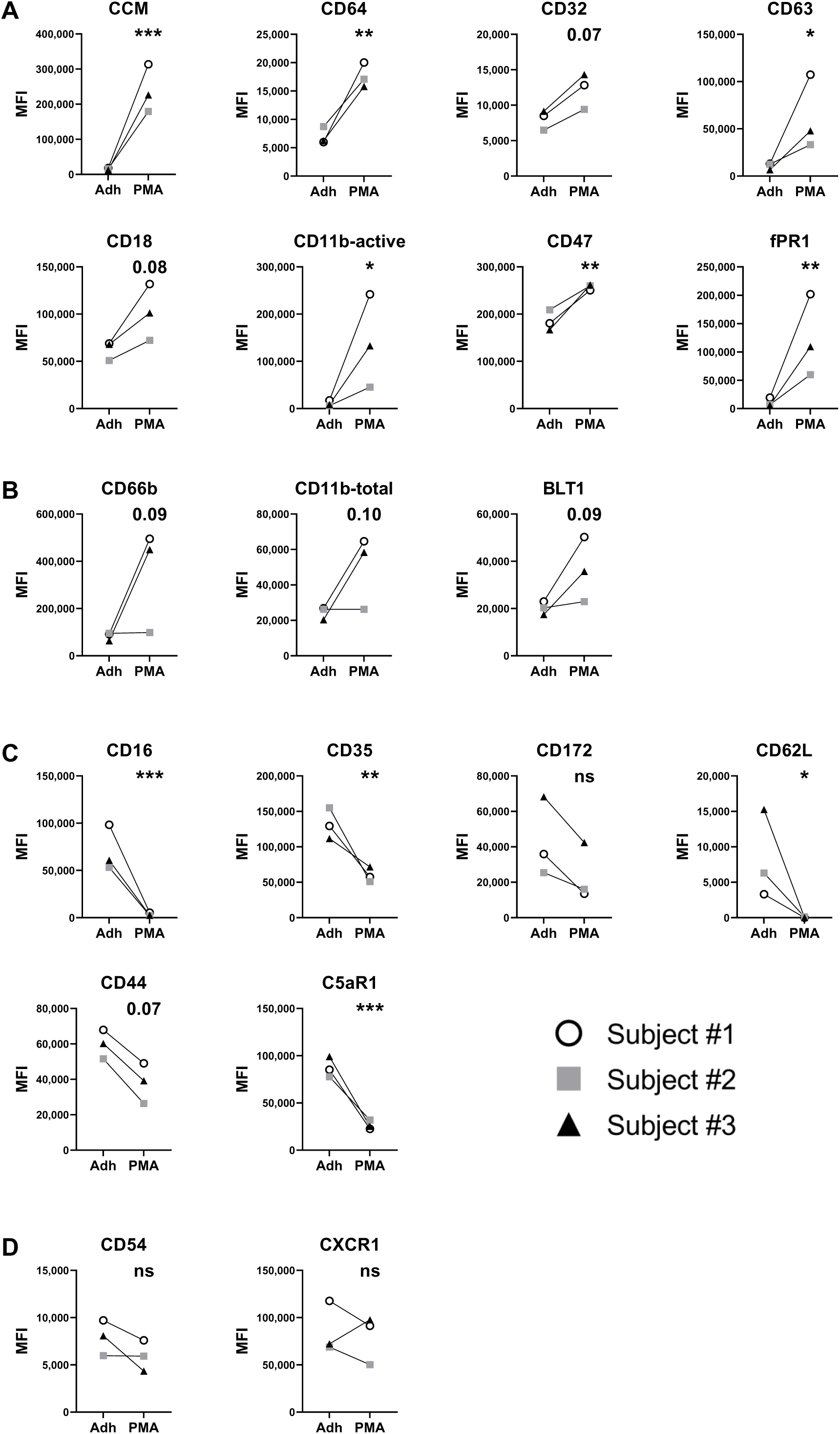
Assessment of Inter-subject Variability of PMN Activation with High-dimensional Spectral Flow Cytometry. Three individual subjects were assayed on separate days. Subject #1 (open circle) was assayed three separate times as in Figure 4 with each replicate averaged together. Subjects #2 (gray square) and #3 (black triangle) were each assayed separately. Surface markers were analyzed via median fluorescence intensity (MFI) for those that were **(A)** consistently upregulated with PMA compared to adherence-alone (Adh) for each subject, **(B)** upregulated in two out of three subjects, **(C)** downregulated in each of the three subjects, or **(D)** showed no consistent trend between the three subjects with PMA stimulation. Statistical analyses were performed on unpaired log-transformed data, Student’s *t*-test. P-value indicated (0.05<p<0.10) or *=p<0.05, **=p<0.01, ***=p<0.001, ****=p<0.0001, ns = not significant. Negative MFI values at the population level were set to a value of 1.0 for graphing, log transformation, and subsequent statistical analyses.

To highlight the high-dimensional and single cell power of spectral flow cytometry we analyzed the adhered and PMA-stimulated PMN data via uniform manifold approximation and projection (UMAP) dimensional reduction.^42^ Cells from the spectral flow data sets were grouped by the UMAP algorithms based on the similarity of their full surface marker repertoires. Shown in Fig. 6 is a representative replicate from Subject #1 in which adherent-alone PMNs and PMA-stimulated PMNs were analyzed via UMAP. PMA-treated PMNs are plotted by individual surface marker intensity, from which surface marker expression patterns and overlap can be qualitatively identified. Additional replicates from Subject #1 and those from Subjects #2 and #3 can be found in Supplemental Figures 2-5.

**Figure 6.**
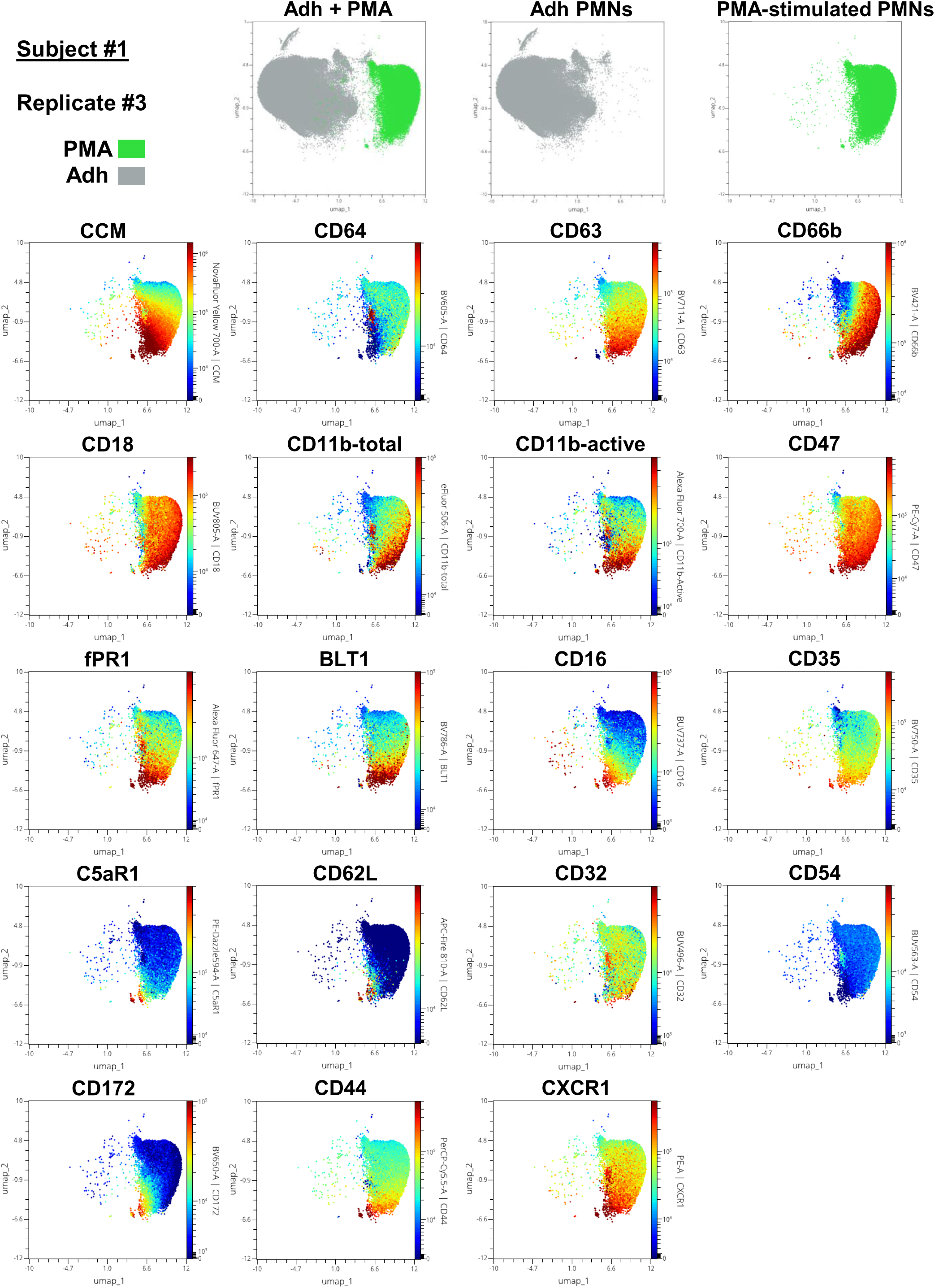
Stimulated PMN Phenotypic Subsets Following UMAP Dimensional Reduction. A representative replicate of Subject #1’s PMA-stimulated PMNs and stained with the full spectral flow cytometry panel were processed and analyzed via uniform manifold approximation and projection (UMAP) dimensional reduction using OMIQ flow cytometry software. Top row: adherent-alone (Adh, gray) and PMA-stimulated (PMA, green) PMN events arrayed by the two principal UMAP components (umap_1 and umap_2). Each of the 19 surface markers analyzed on live PMNs is displayed by the two principal UMAP components and by color intensity (red = highest surface expression, blue = lowest surface expression).

Taken together, these results demonstrate that trends in PMN responses to a known activating stimulus can be identified using this multiparametric panel and that multidimensional analysis can identify unique populations for subsequent investigation.

### PMN responsiveness to the bacterial pathogen Neisseria gonorrhoeae

To demonstrate the utility of this panel to interrogate PMN interactions with phagocytic cargo, including pathogens, CellTrace Blue-labeled *N. gonorrhoeae* was introduced to adherent PMNs at a ratio of either 1 or 10 bacteria per PMN (multiplicity of infection, MOI). A single subject’s PMNs were challenged on three independent days with fluorescent bacteria, unlabeled control bacteria, or adherent-alone PMNs with no bacteria exposure.

Following 1hr of infection, eight PMN markers showed consistently increased trends in surface expression with *N. gonorrhoeae* challenge, which increased with MOI (CCM, CD64, CD63, CD66b, fPR1, CD18, CD11b-active, and BLT1; Fig. 7A). Three other markers, CD11b-total, CD35, and CD47, had increased surface expression with infection at an MOI of 1 but a decrease at an MOI of 10; the MFI of CD35 was lower at MOI of 10 than in the uninfected population (Fig. 7B). Five markers (CD16, CD172, CD62L, C5aR1, and CD44) consistently decreased in surface expression after infection with *N. gonorrhoeae* (Fig. 7C). The remaining 3/19 PMN markers had no consistent trends between the three experimental replicates (Fig. 7D).

**Figure 7.**
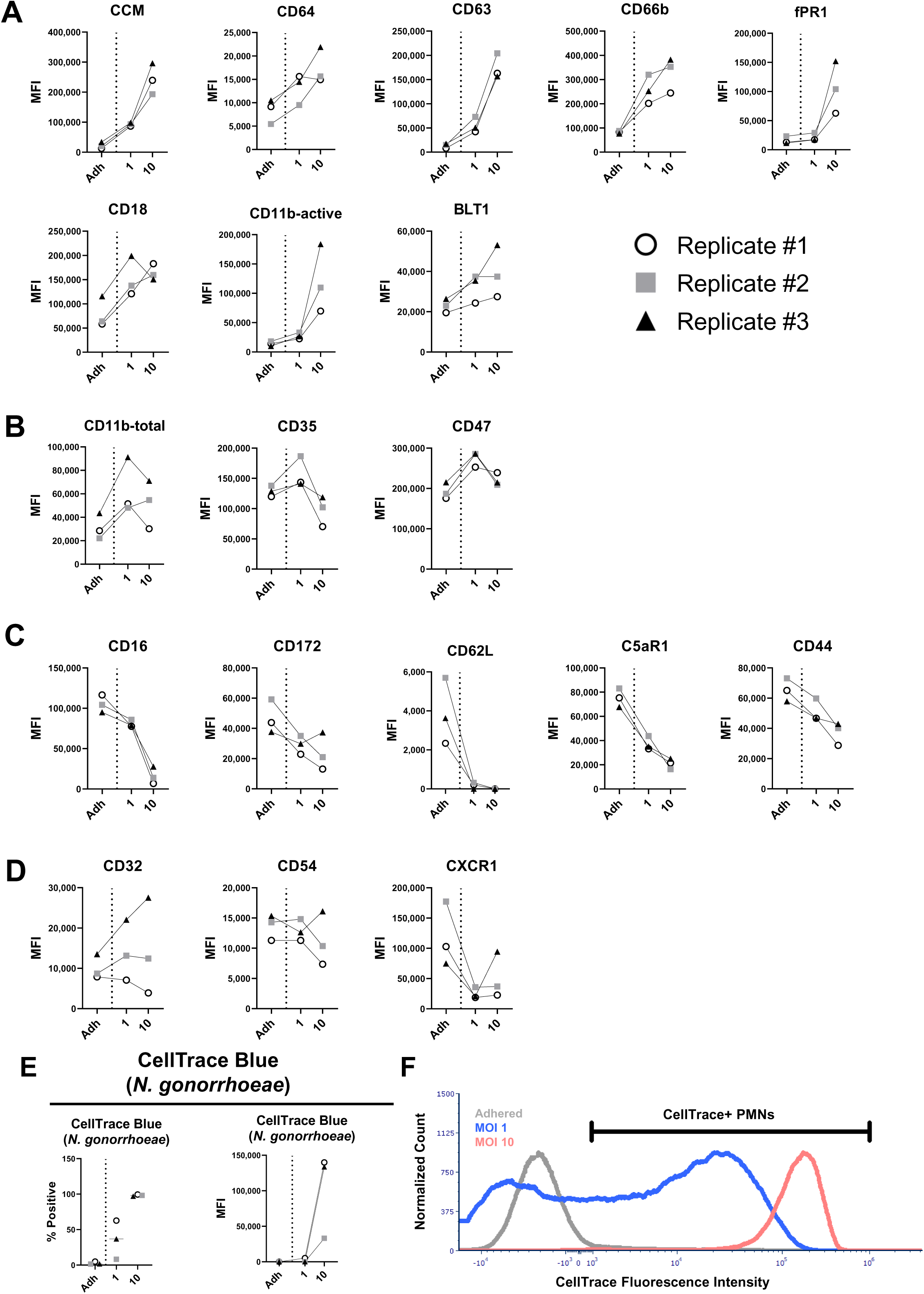
Fluorescently-labeled Bacteria Alter PMN Surface Protein Expression in an Infectious Dose-Dependent Manner. Primary human PMNs were adhered as above with or without challenge with CellTrace Blue-labeled *N. gonorrhoeae* at a multiplicity of infection (MOI) of 1 or 10 bacteria per PMN and analyzed via spectral flow cytometry. **(A)** Surface proteins which were consistently elevated upon *Neisseria* infection compared to adherent-alone (Adh) conditions and trended upwards in a dose-dependent manner (MOI 1 versus 10). **(B)** PMN markers which increased with an infection of 1 bacterium per PMN over adherent-alone PMNs but decreased in the MOI of 10 condition versus MOI of 1. **(C)** Surface markers which consistently decreased with bacterial infection in each replicate in an infectious dose-dependent manner. **(D)** Surface markers which showed no consistent trends between replicates of *N. gonorrhoeae* infection. **(E)** Percent positivity and CellTrace Blue MFI of PMNs in each condition as a representation of direct association with fluorescently-labeled *N. gonorrhoeae.* **(F)** Representative distribution of CellTrace Blue intensity in the adherent alone, MOI 1, and MOI 10 conditions. Marker denotes gating limits of positive CellTrace Blue signal.

We next examined CellTrace Blue fluorescence as a correlate of bacterial burden encountered by PMNs within each population. PMNs were gated into CellTrace Blue-positive and negative populations. PMNs infected at an MOI of 1 exhibited 8.2 to 62.7 percent CellTrace Blue positivity, while PMNs at an MOI of 10 were greater than 97 percent positive in all biological replicates (Fig. 7E). The measured MFI for PMNs infected at an MOI of 10 was more than a log_10_-fold greater than those in the MOI of 1, and the population distribution of fluorescence intensity was more homogenous (coefficient of variance 49.4 versus 104.9; Fig. 7F,G). The observed CellTrace Blue heterogeneity in the MOI of 1 prompted more nuanced investigation into how PMN surface markers varied with bacterial burdens.

To verify that CellTrace intensity corresponded with bacterial burden, PMNs exposed to *N. gonorrhoeae* at an MOI of 1 were examined using imaging flow cytometry in which single, focused cells were gated by CellTrace MFI into CellTrace-negative PMNs with ‘No *Neisseria*’, the lowest quartile of CellTrace-positive PMNs as the ‘Low *Neisseria*’ population, and the highest quartile of CellTrace-positive PMNs as the ‘High *Neisseria*’ population (Fig. 8A). The No *Neisseria* gate represented 20-50% of the total population (Fig. 8B). The High *Neisseria* population’s MFI was 19.2-fold greater than the Low population’s MFI; there was negligible fluorescence in the No *Neisseria* population (Fig. 8B). Bacterial burden was verified by imaging flow cytometry as CellTrace intensity corresponded with the numbers of bacteria directly associated with individual PMNs which also showed that PMNs in the High *Neisseria* gate frequently contained 10 or more bacteria (Fig. 8C).

**Figure 8.**
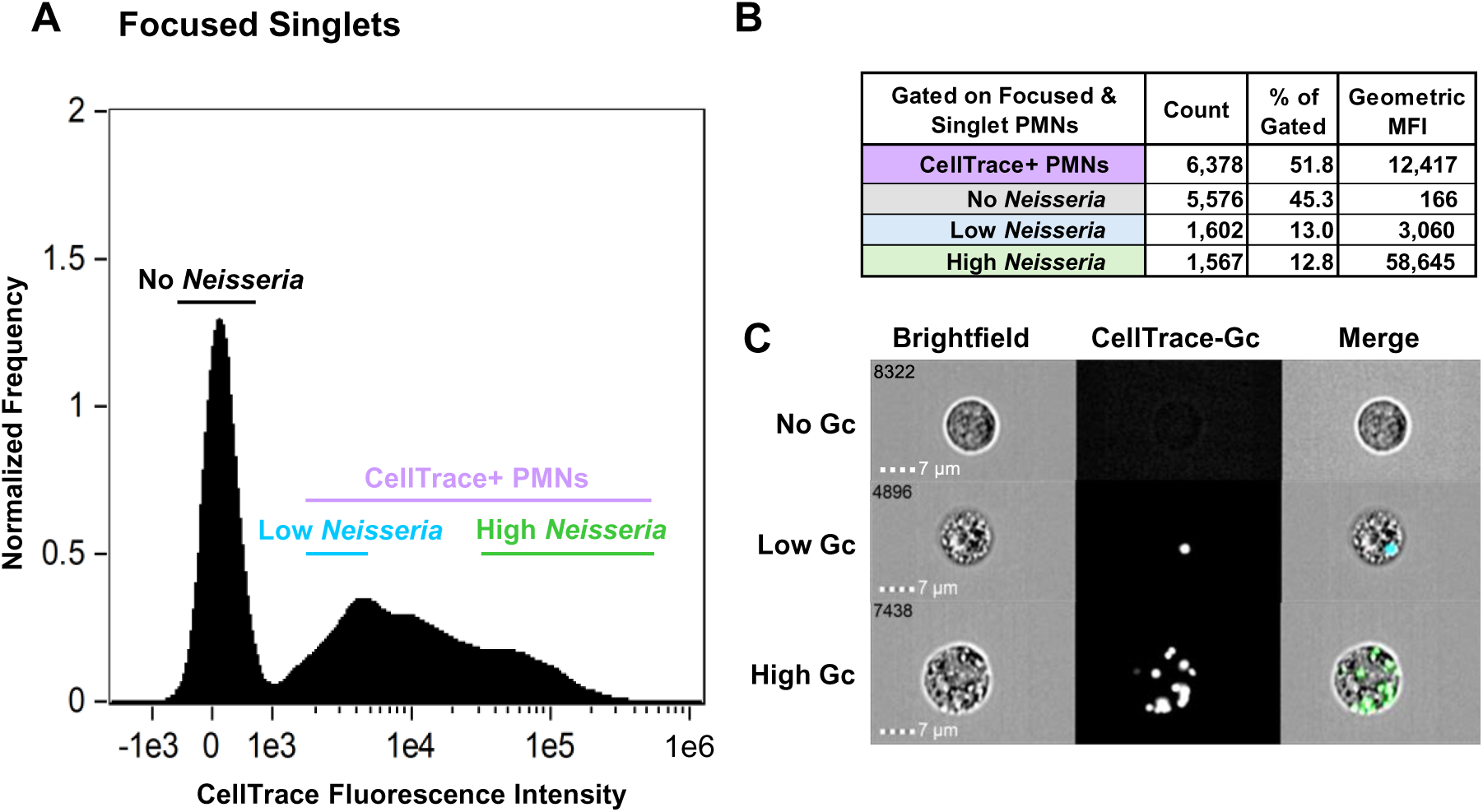
Imaging Flow Cytometry Demonstrates Bacterial Burden on Single PMN Level. Primary human PMNs were isolated and adhered as described and infected with CellTrace-labeled *N. gonorrhoeae* at a ratio of 1 bacterium per PMN for 1 hour and assayed via imaging flow cytometry. **(A)** Focused, single cell events were gated into events based on CellTrace median fluorescence intensity. CellTrace-negative events were classified as having No *N. gonorrhoeae*, whereas CellTrace-positive PMNs were classified as being associated with *N. gonorrhoeae*. CellTrace-positive PMNs were further subdivided into *N. gonorrhoeae* Low and *N. gonorrhoeae* High groups by quartile. **(B)** Statistics from panel (A) are shown with number of events, percentages, and geometric median fluorescent intensities (MFIs) in each gate. **(C)** Representative micrographs from each gate of panel (A) showing the brightfield channel, CellTrace/*N. gonorrhoeae* channel and a merged channel demonstrating bacterial burden on a single PMN basis. Numbers in the top lefthand corner of each image series represents the event number acquired out of ten thousand individual focused singlet events.

Using spectral flow cytometry, MOI of 1 PMNs (see Fig. 7) were subgated into No, Low, and High *Neisseria* populations as above; also included were adherent-alone PMNs which had not been exposed to bacteria (Fig. 9A). The CellTrace Blue MFI for the uninfected and No *Neisseria* populations were undetectable, whereas the High *Neisseria* population had a 9.8-fold greater MFI on average than the Low *Neisseria* population (Fig. 9A,B). Taken together, the imaging and spectral flow cytometry demonstrate that the MOI of 1 experimental condition yields a population of PMNs that exhibit a range of interactions with bacteria, despite exposure to the same inoculum of infectious particles.

**Figure 9.**
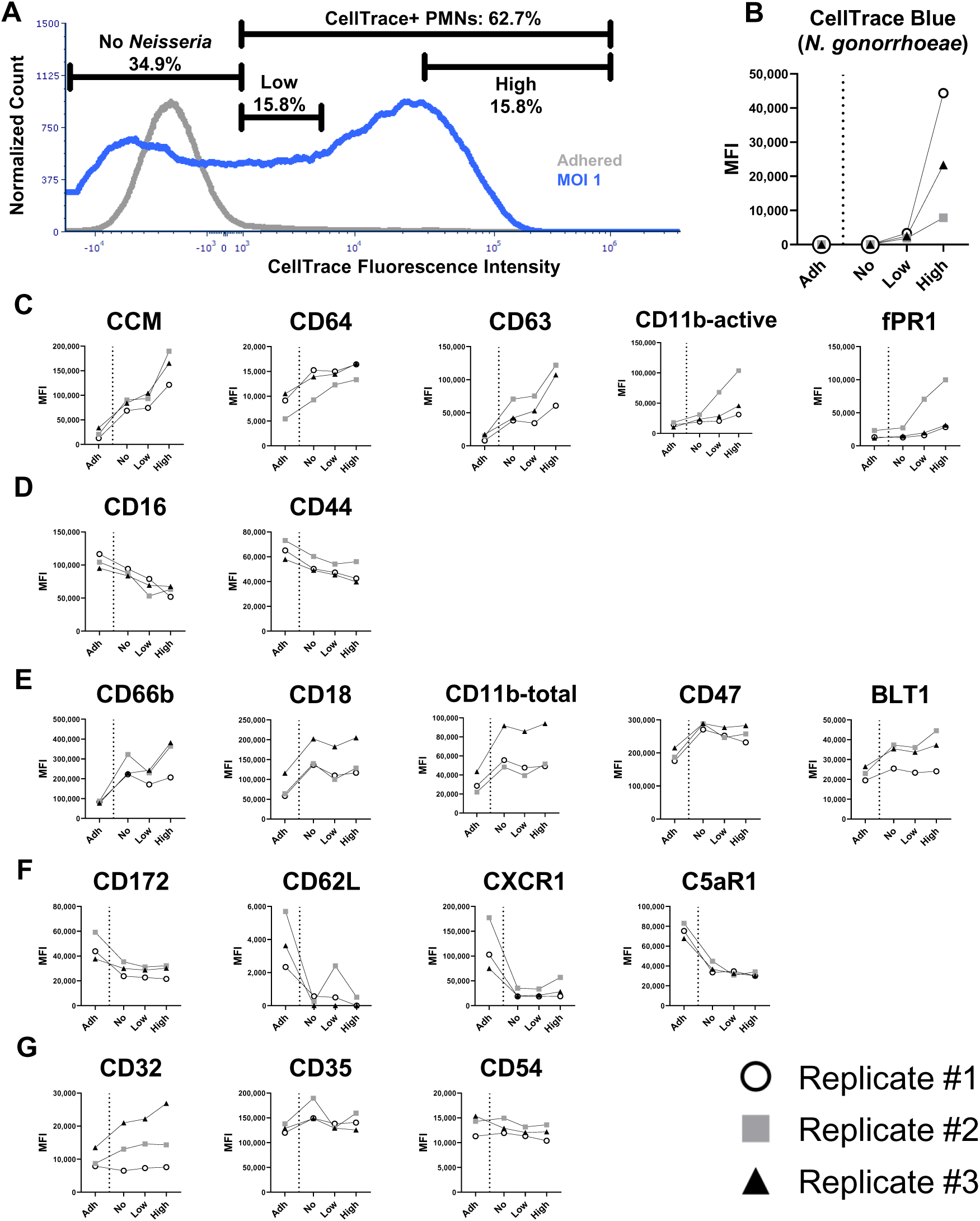
Differential Surface Marker Expression Patterns Based on Bacterial Burden and Direct Association. **(A)** Histogram of CellTrace positivity from Figure 8 which were challenged with CellTrace-Blue labeled *N. gonorrhoeae* at a multiplicity of infection (MOI) of 1 bacterium per PMN or adhered alone without bacterial exposure (Adh). PMNs in the *N. gonorrhoeae* infected condition were subgated by CellTrace Blue intensity as described in Figure 8 into a No *Neisseria* population and populations with Low and High bacterial burdens based on fluorescence quartiles. **(B)** CellTrace Blue/*N. gonorrhoeae* MFI from three independent experiments (different colors/shapes) separated by bacterial burden subgate. Adherent-alone PMNs (Adh) to the left of the dotted line displayed as a control. **(C)** PMN markers which consistently increased with *N. gonorrhoeae* infection over adhered-alone conditions and in a bacterial burden-dependent manner, i.e. High *Neisseria* over Low *Neisseria*. **(D)** Surface markers which consistently decreased in a bacteria burden-dependent manner and compared to adherent-alone. **(E)** Surface markers which consistently increased with *N. gonorrhoeae* infection compared to adhered-alone PMNs, but with no bacteria burden-dependent variation in surface protein expression. **(F)** Surface markers which consistently decreased with *N. gonorrhoeae* infection compared to adhered alone PMNs but with no bacterial burden-dependent variation in surface protein expression. **(G)** PMN surface markers which showed either no change in expression or no consistent trends in variation between replicates.

The No, Low, and High *Neisseria* populations were examined for surface markers expression using the spectral flow cytometry panel and compared with the uninfected control PMNs. The MFIs of CCM, CD64, CD63, CD11b-active, and fPR1 all increased in a bacterial burden-dependent manner (Fig. 9C). Interestingly, CCM, CD64, and CD63 surface expression were greater in the No *Neisseria* population from *N. gonorrhoeae* exposed PMNs than in the uninfected controls (Fig. 9C). Conversely, CD16 and CD44 decreased consistently in a bacterial burden-dependent manner, and the uninfected control had a higher MFI than the No *Neisseria* population from the bacteria exposed PMNs (Fig. 9D).

Other markers changed in surface expression levels with *N. gonorrhoeae* infection compared to the uninfected control, but did not vary with MOI across the No, Low, and High *Neisseria* populations: CD66b, CD18, CD11b-total, CD47, and BLT1 all increased in the presence of *N. gonorrhoeae* (Fig. 9E), whereas CD172, CD62L, CXCR1, and C5aR1 all decreased (Fig. 9F). CD32, CD35, and CD54 did not consistently respond to infection across replicates or showed no change regardless of bacterial burden (Fig. 9G).

## Discussion

PMNs have major roles in infection, sterile inflammation, and tissue injury and repair. Here we describe the development, validation, and application of a high-dimensional spectral flow cytometry panel for profiling PMNs isolated from human subjects. It was designed to interrogate PMN activation and surface expression of markers for cellular functionality including phagocytosis, degranulation, migration, and chemotaxis. This spectral flow cytometry panel enables rapid, high throughput, multiparametric analysis, and sample preservation with broad applicability to PMN biology. As highlighted in the panel design section (Fig. 2), care was taken to allow adaptability of this panel. For example, in place of CCMs as surface receptors that bind and phagocytose *N. gonorrhoeae* in the NFY700 channel, a research team could substitute a marker or receptor of their interest. In the CTB/BUV395 channel, another surface marker or labeled cargo could be studied such as other pathogens, synthetic particles with different chemical/physical properties, cellular debris, or immune complexes. Additional PMN functions that could be incorporated into the panel include ROS generation, NET release, death modalities including apoptosis or pyroptosis, and/or intracellular signaling/cytokine production.

As highly differentiated, terminal cells, PMNs present unique challenges for building and applying a flow cytometric panel. Measuring PMN activation between a baseline and stimulated state is inherently challenging due PMNs’ sensitivity to activation including during the isolation procedure. Here, PMNs were isolated by Ficoll gradient and hypotonic erythrocyte lysis, which differ in basal activation state from PMNs in anticoagulated whole blood, purified by immunomagnetic negative selection, or in tissues.^61^ The surface epitope density of a selected marker can vary dramatically on a continuum across resting, primed, and activated states. Furthermore, some surface markers are shed or internalized by activated PMNs. For these reasons, the selection of marker-fluorochrome pairings, which is based on epitope density and brightness index, must be experimentally determined through antibody titrations in single-stained samples at low and high epitope density conditions as well as in the context of the full stained panel. Fluorescence-minus-one (FMO) controls should also be employed where necessary to discriminate between positive and negative populations for accurate gate setting.^62,63^ Finally, there is PMN surface marker variability basally and upon stimulation for single subject on different days and between subjects (Fig. 4 and 5). Given the day-to-day variability observed within a single individual’s PMN responses, it may be prudent to assess subjects multiple times. Future studies are needed to define the number of individuals or experiments on a single subject needed for statistical power.

PMNs play a preeminent role in controlling pathogenic organisms. However, some pathogens have evolved to counteract the antimicrobial mechanisms of PMNs. Among these is the Gram-negative bacterium *Neisseria gonorrhoeae* which infects mucosal surfaces of its obligate human host and stimulates a PMN-driven inflammatory response.^64,65^ Here, *N. gonorrhoeae* was chosen as a model infectious organism and phagocytic cargo for its capacity to survive within phagosomes of human neutrophils.^66–68^ By engaging neutrophil surface receptors *N. gonorrhoeae* can block neutrophil phagosome maturation and suppress neutrophil activation, such that viable bacteria are isolated from the PMN-rich exudates of infected individuals.^69–71^ Intriguingly, PMNs in urethral gonorrheal exudates have a heterogenous distribution of associated *N. gonorrhoeae* resembling that of Fig. 8 in which many PMNs have no bacteria, some having single-digit numbers of bacteria, and others having tens of bacteria.^72^ We anticipate that spectral flow cytometry can be applied to understand the biology underlying this diversity. During infection, PMNs interact directly with in-tact microbes as well as the soluble factors they release such as cell wall fragments, formylated peptides, and lipo-poly/oligo-saccharides, alone or in outer membrane vesicles.^7^ Intriguingly, we observed differences in surface marker expression between PMNs not exposed to *N. gonorrhoeae* and those in the infection milieu that did not contain associated or phagocytosed bacteria (Fig. 8,9). This observation implies that soluble factors within the infectious milieu are sufficient to initiate PMN activation, which is enhanced with bacterial association (i.e. in the Low/High *Neisseria* conditions). Additionally, paracrine signaling among PMNs during infection, such as release of leukotriene-B_4_ could contribute to the observed changes in some surface markers.^73^ Our findings prompt further investigation with purified soluble factors and/or outer membrane vesicles and *N. gonorrhoeae* mutants that produce or release different quantities of these factors.

PMNs sense many stimuli in the context of infection, tissue damage, or inflammation and integrate these signals to respond. The field now appreciates that PMNs vary within a population by their age and prior experience (“trained innate immunity”),^74^ yet the contribution of this diversity in host response is incompletely understood. We developed this high-dimensional spectral flow cytometry panel to profile the diversity of PMN responses, identify subsets of PMNs with phenotypes of interest to the investigator, and generate new hypotheses about human PMN functionality. Application of high-dimensional analytics (Fig. 6) offers a significant ability to decipher underlying biology from the vast amounts of data generated from this panel. Going forward, this panel can be directly transferred to spectral cell sorting instruments with appropriate laser and detector configurations in order to isolate PMN populations of interest. This panel and protocol is also suitable for collection of viable PMNs containing *N. gonorrhoeae* which can be used for downstream applications such as single cell RNA-sequencing (Supplemental Fig. 6).^75,76^ As spectral flow cytometers and the technology associated with them continue to advance, the human PMN functional panel presented here can be used and expanded by many investigators to understand the breadth of responses of PMNs, uncovering the contribution of these biologically meaningful cells to diverse fields of biology and medicine.

## Acknowledgements and Sources of Funding

We thank members of the Criss Lab and UVA Flow Cytometry Core Facility (FCCF), past and present, for their advice and insights. We also thank Dr. Loren Erickson (UVA) for his input and guidance on multiparametric spectral flow cytometry analysis. This work was supported by NIH R01 AI097312, U19 AI158930, U19 AI144180, and U01 AI162457. The UVA FCCF (RRid:SCR_017829) was supported in part by NCI P30 CA044579. ERL was supported in part by NIH F30 AI179038, T32 AI007046, and T32 GM007267.

## Authorship Contribution Statement

<u>Evan R. Lamb:</u> Conceptualization, Methodology, Analysis, Investigation, Writing – Original Draft, Validation, Visualization. <u>Ian J. Glomski:</u> Analysis, Investigation, Writing – Original Draft, Validation, Visualization. <u>Taylor A. Harper:</u> Methodology, Analysis, Investigation, Writing – Review & Editing. <u>Michael D. Solga:</u> Methodology, Analysis, Writing – Review & Editing, Funding Acquisition. <u>Alison K. Criss:</u> Conceptualization, Methodology, Analysis, Writing – Original Draft, Writing – Review & Editing, Funding Acquisition, Project Administration, Supervision

---

**SUPPLEMENTAL FIGURE 1.** Ficoll-purified cells were adhered and stained (see Materials and Methods) for markers enriched in different leukocyte populations. **(A)** Cells were gated into PMNs based on the characteristic forward and side scatter profiles (FSC-A, SSC-A), single cells events, and live cells by Zombie near-infrared (NIR) exclusion. **(B)** Cells were gated by positivity for canonical neutrophilic markers CD16, CD66b, and CD11b. Cells were further assessed for their low expression of the major monocyte surface marker CD14 **(C)** or eosinophil marker CD49d **(D)**.

**SUPPLEMENTAL FIGURE 2.** Purified primary human PMNs from Subject #1 replicate #1 that were stimulated or not with PMA were stained with the full spectral flow cytometry panel. PMNs were then processed and analyzed via UMAP dimensional reduction using OMIQ flow cytometry software. Top row: adherent-alone (Adh, gray) and PMA-stimulated (PMA, red) PMN events arrayed by the two principal UMAP components (umap_1 and umap_2). Each of the 19 surface markers analyzed on live, PMA-treated PMNs is displayed by the two principal UMAP components and by color intensity (red = highest surface expression, blue = lowest surface expression).

**SUPPLEMENTAL FIGURE 3.** UMAP dimensional reduction of the spectral flow cytometry profiles of PMNs from Subject #1, replicate #2, with and without PMA treatment, conducted as in Supplemental Figure 2. Top row: adherent-alone (Adh, gray) and PMA-stimulated (PMA, cyan) PMN events arrayed by the two principal UMAP components (umap_1 and umap_2). Each of the 19 surface markers analyzed on live, PMA-treated PMNs is displayed by the two principal UMAP components and by color intensity (red = highest surface expression, blue = lowest surface expression).

**SUPPLEMENTAL FIGURE 4.** UMAP dimensional reduction of the spectral flow cytometry profiles of PMNs from Subject #2, replicate #1, with and without PMA treatment, conducted as in Supplemental Figure 2. Top row: adherent-alone (Adh, gray) and PMA-stimulated (PMA, violet) PMN events arrayed by the two principal UMAP components (umap_1 and umap_2). Each of the 19 surface markers analyzed on live, PMA-treated PMNs is displayed by the two principal UMAP components and by color intensity (red = highest surface expression, blue = lowest surface expression).

**SUPPLEMENTAL FIGURE 5.** UMAP dimensional reduction of the spectral flow cytometry profiles of PMNs from Subject #3, replicate #1, with and without PMA treatment, conducted as in Supplemental Figure 2. Top row: adherent-alone (Adh, gray) and PMA-stimulated (PMA, blue) PMN events arrayed by the two principal UMAP components (umap_1 and umap_2). Each of the 19 surface markers analyzed on live, PMA-treated PMNs is displayed by the two principal UMAP components and by color intensity (red = highest surface expression, blue = lowest surface expression).

**SUPPLEMENTAL FIGURE 6.** Adherent primary human PMNs were infected with CellTrace-labeled *N. gonorrhoeae* at a ratio of 1 bacterium per PMN for 1 hr then collected for spectral cell sorting flow cytometry as described. PMNs were stained with Zombie near-infrared (NIR) viability dye and run on the Cytek Aurora CS Cell Sorter, with the PMNs collected into Eppendorf tubes. Shown is the distribution of ZNIR viability dye fluorescence intensity from post-sorting PMNs (violet) compared to a mixture of live and heat-killed PMNs (gray). Y-axis reports the normalized count frequency for the number of collected events.

