## Supplemental Figures and legends for "High-dimensional spectral flow cytometry of activation and phagocytosis by peripheral human polymorphonuclear leukocytes"

**SUPPLEMENTAL FIGURE 6.** Adherent primary human PMNs were infected with CellTrace-labeled *N. gonorrhoeae* at a ratio of 1 bacterium per PMN for 1 hr then collected for spectral cell sorting flow cytometry as described. PMNs were stained with Zombie near-infrared (NIR) viability dye and run on the Cytek Aurora CS Cell Sorter, with the PMNs collected into Eppendorf tubes. Shown is the distribution of ZNIR viability dye fluorescence intensity from post-sorting PMNs (violet) compared to a mixture of live and heat-killed PMNs (gray). Y-axis reports the normalized count frequency for the number of collected events.

#### Supplemental Figure 1

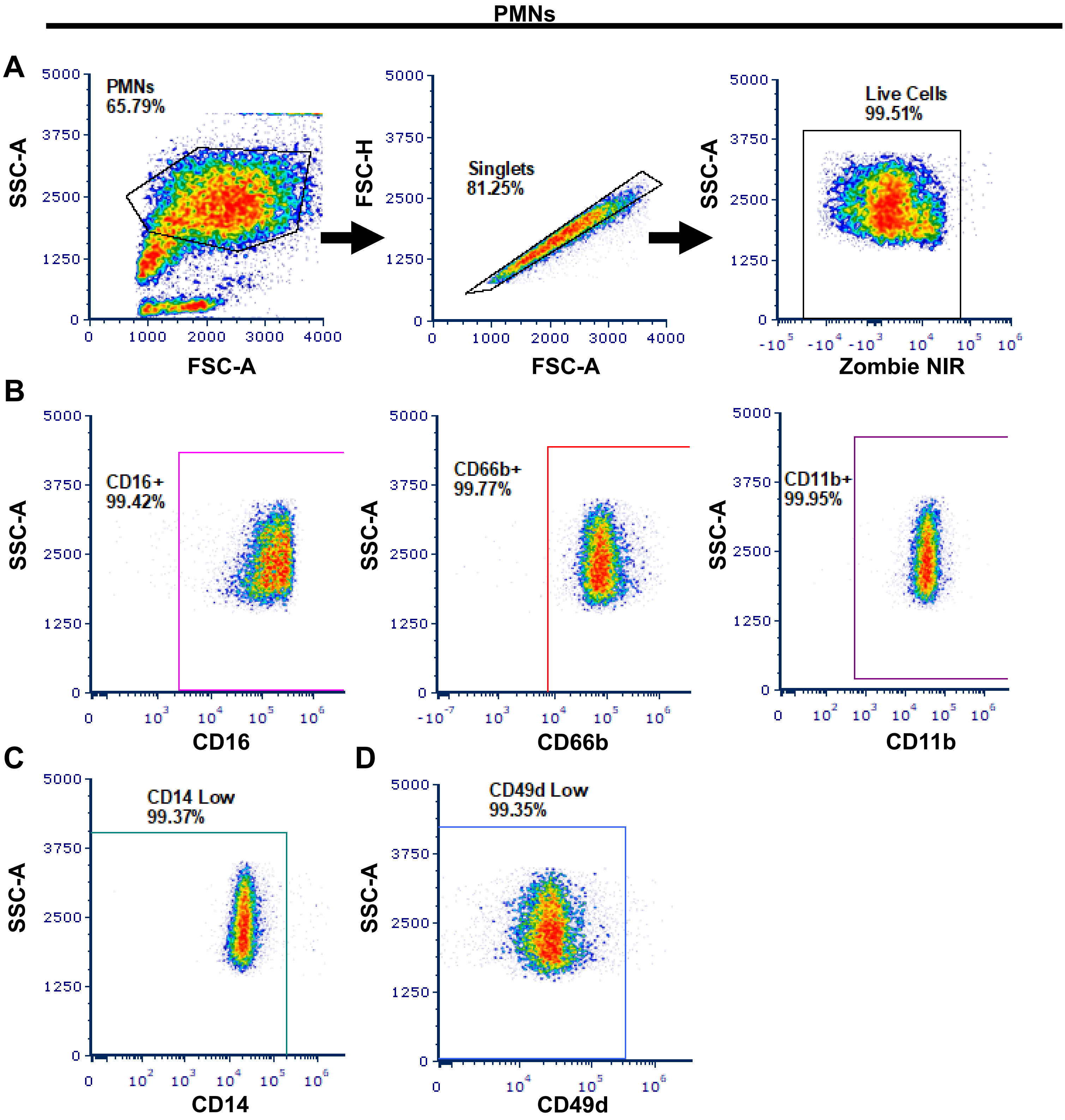

### Supplemental Figure 2

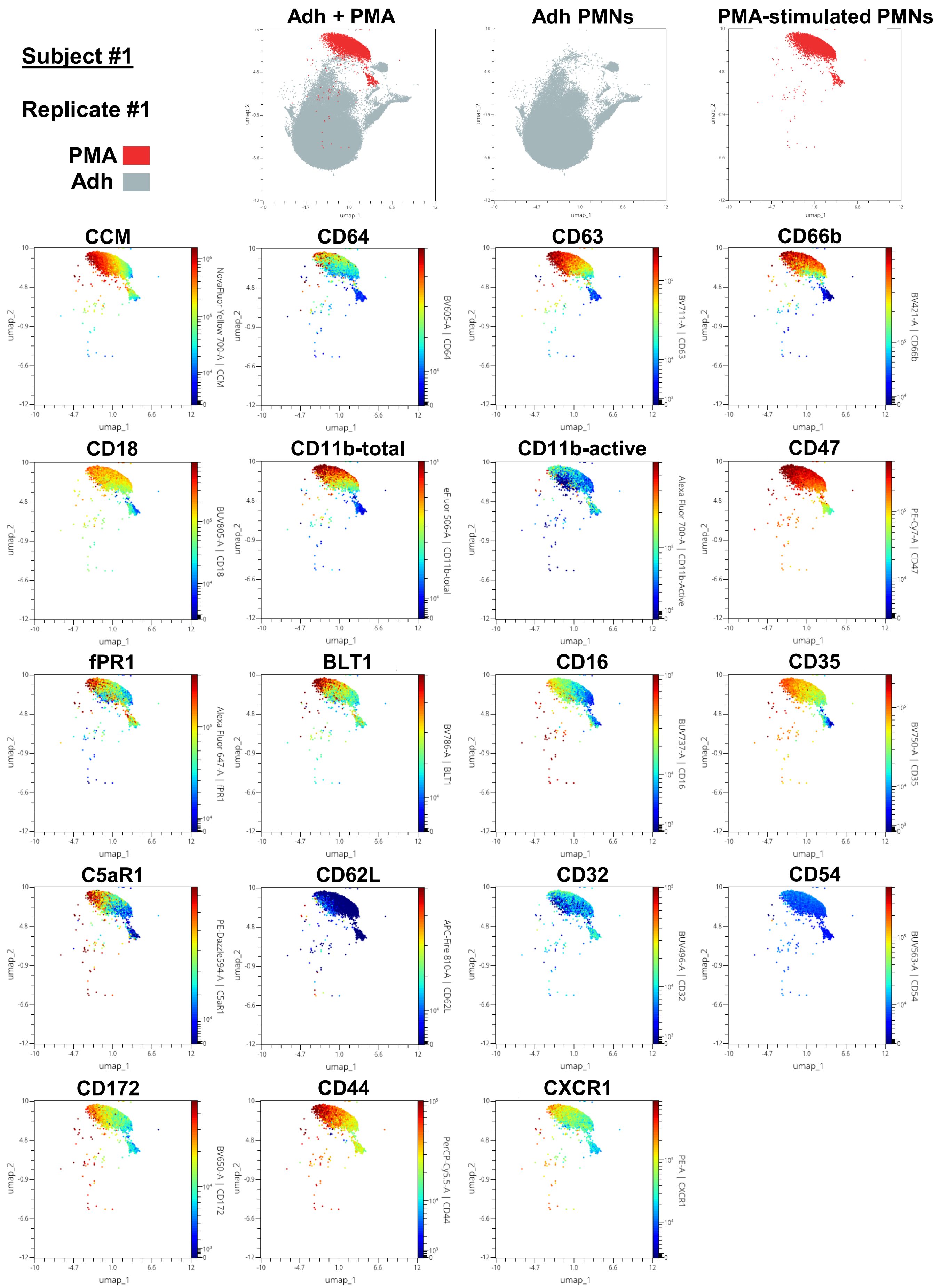

### Supplemental Figure 3

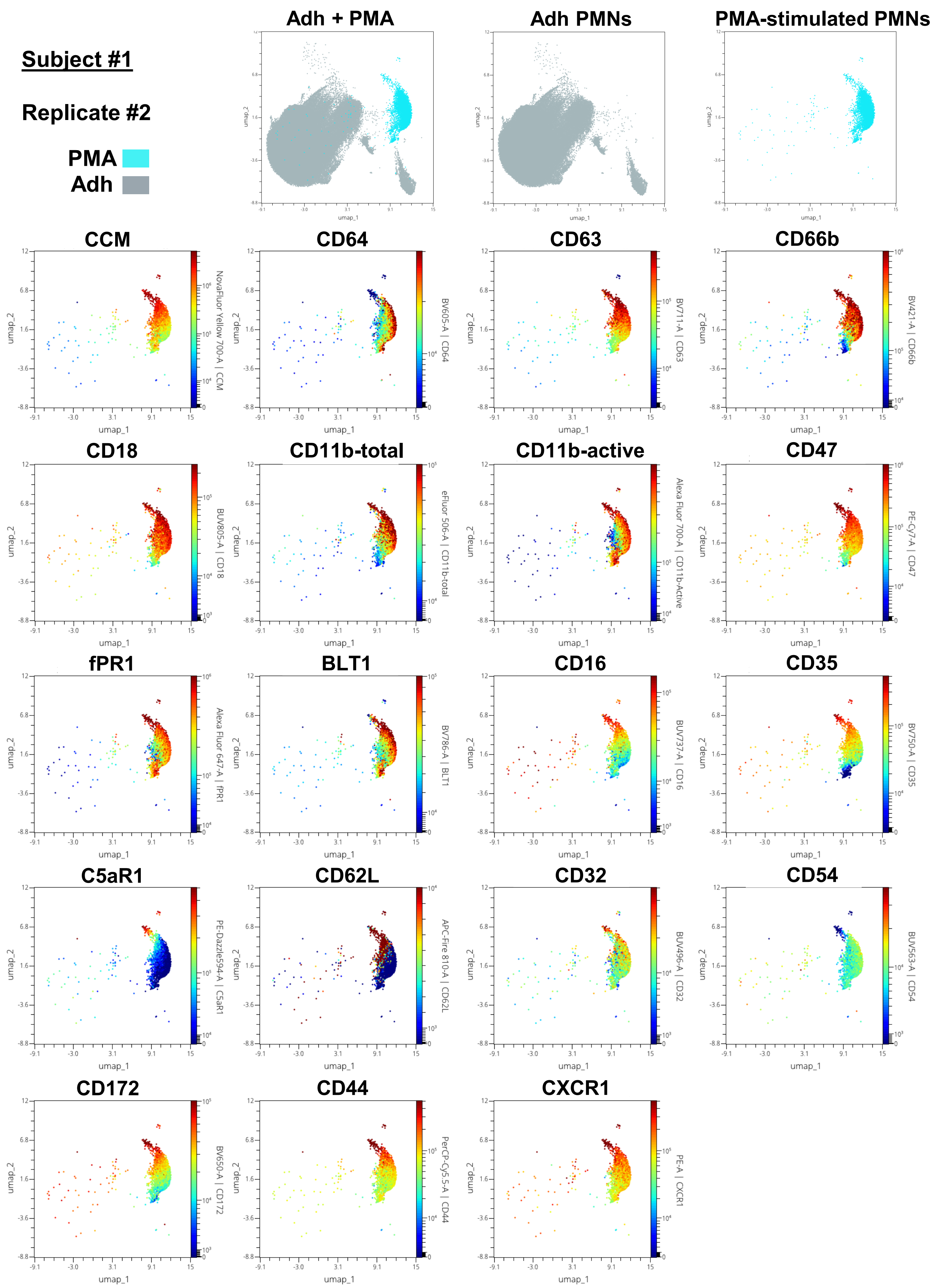

### Supplemental Figure 4

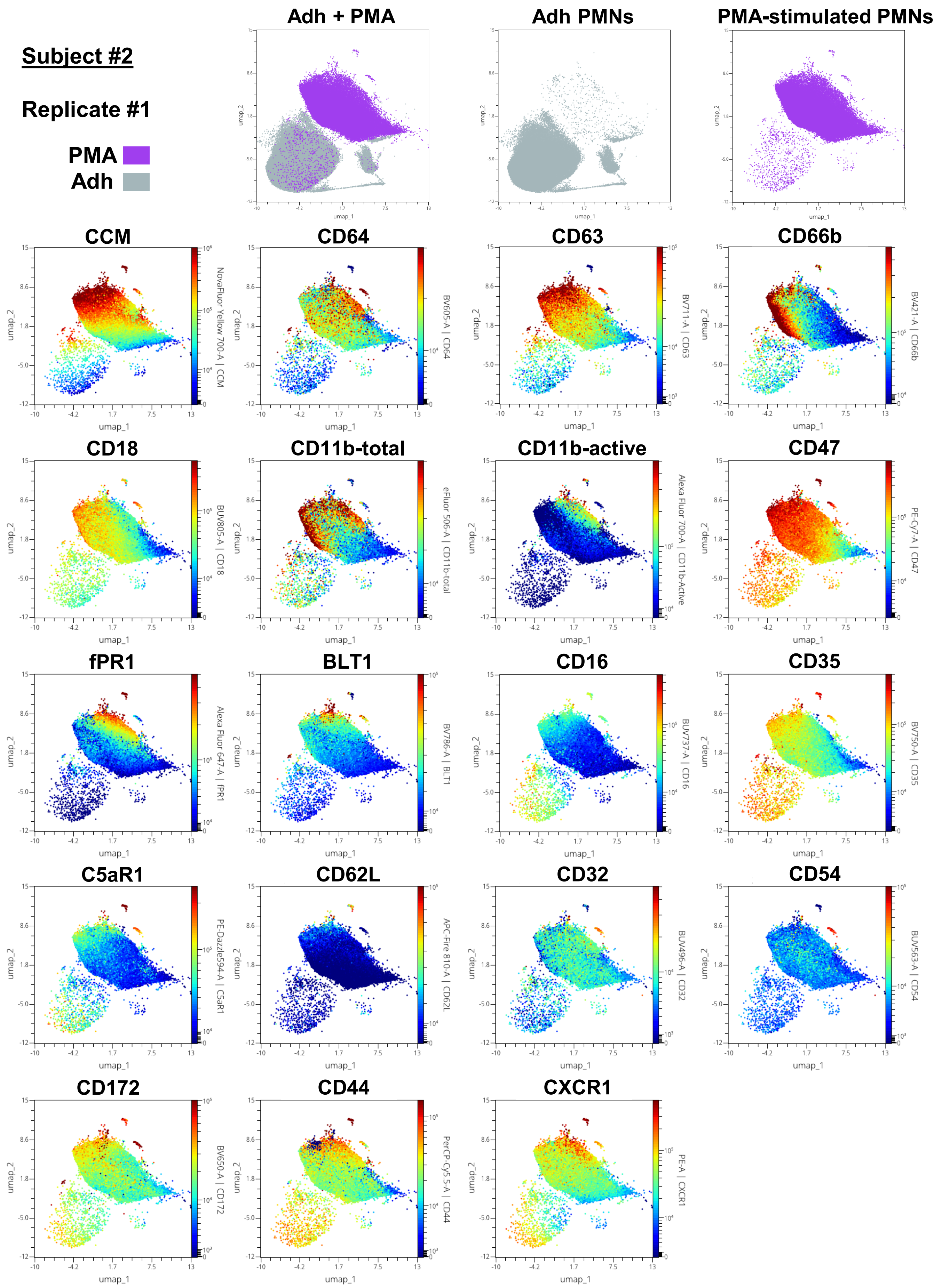

### Supplemental Figure 5

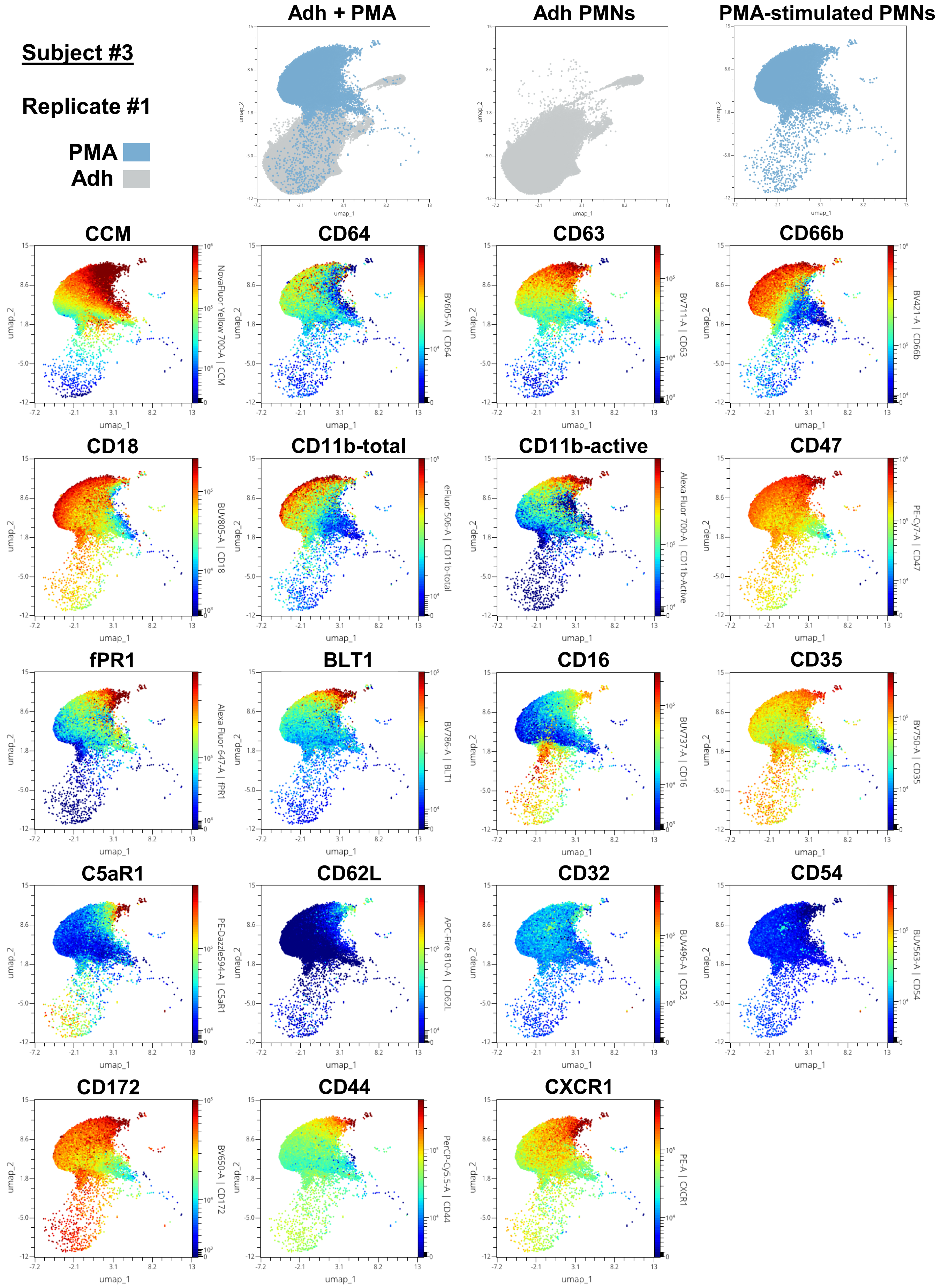

**Supplemental Figure 6**

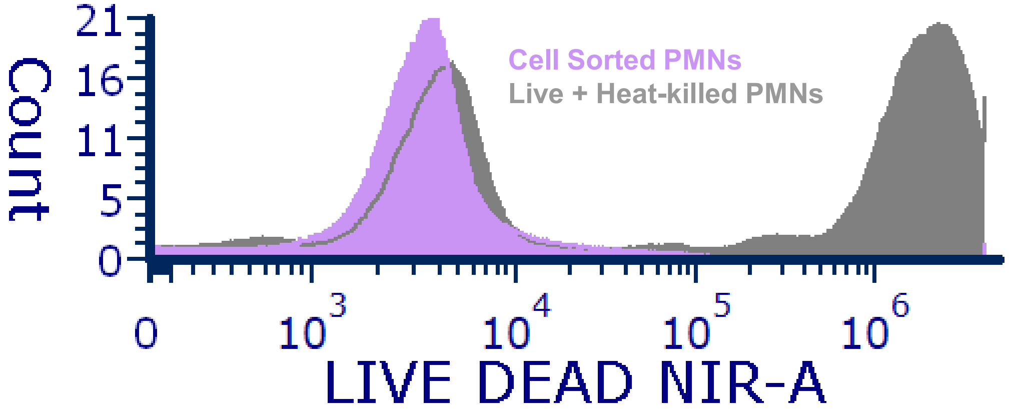
